# Discovery of Potent Triple Inhibitors of Both SARS-CoV-2 Proteases and Human Cathepsin L

**DOI:** 10.1101/2021.10.19.465036

**Authors:** Ittipat Meewan, Jacob Kattoula, Julius Y. Kattoula, Danielle Skinner, Pavla Fajtová, Miriam A. Giardini, Brendon Woodworth, James H. McKerrow, Jair Lage de Siqueira-Neto, Anthony J. O’Donoghue, Ruben Abagyan

**Affiliations:** Skaggs School of Pharmacy and Pharmaceutical Sciences University of California San Diego, La Jolla, California 92093, United States; Department of Chemistry and Biochemistry University of California San Diego, La Jolla, California 92093, United States; Biological Sciences University of California San Diego, La Jolla, California 92093, United States

**Keywords:** Covid19 drug candidates, Multiple protease inhibitors, Disulfiram, Thiuram disulfide, dithiobis-(thioformate), SARS-CoV-2 Main Protease, SARS-CoV-2 Papain Like Protease, Cathepsin L, Transmembrane Serine Protease 2 TMPRSS2, COVID-19

## Abstract

There are currently no FDA approved inhibitors of SARS-CoV-2 viral proteases with specific treatment for post-exposure of SARS-CoV-2. Here, we discovered inhibitors containing thiuram disulfide or dithiobis-(thioformate) tested against three key proteases in SARS CoV-2 replication including SARS CoV-2 Main Protease (Mpro), SARS CoV-2 Papain Like Protease (PLpro), and human cathepsin L. The use of thiuram disulfide and dithiobis-(thioformate) covalent inhibitor warheads was inspired by disulfiram, a currently prescribed drug commonly used to treat chronic alcoholism that at the present time is in Phase 2 clinical trials against SARS-CoV-2. At the maximal allowed dose, disulfiram is associated with adverse effects. Our goal was to find more potent inhibitors that target both viral proteases and one essential human protease to reduce the dosage and minimize the adverse effects associated with these agents. We found that compounds coded as RI175, JX 06, and RI172 are the most potent inhibitors from an enzymatic assay against SARS-CoV-2 Mpro, SARS-CoV-2 PLpro, and human cathepsin L with IC_50_s of 330, 250 nM, and 190 nM about 4.5, 17, and 11.5-fold more potent than disulfiram, respectively. The identified protease inhibitors in this series were also tested against SARS CoV-2 in a cell-based and toxicity assay and were shown to have similar or greater antiviral effect than disulfiram. The identified triple protease inhibitors and their derivatives are promising candidates for treatment of the Covid-19 virus and related variants.

## Introduction

The Severe acute respiratory syndrome coronavirus (SARS-CoV) and the Middle east respiratory syndrome coronavirus (MERS-CoV) are members of coronaviridae that can cause fatal respiratory diseases and are rapidly transmitted, but their outbreaks were far from pandemic in scale. In December 2019, a new coronavirus known as Severe acute respiratory syndrome coronavirus 2 (SARS-CoV-2) was identified in Wuhan China that causes coronavirus disease 2019 (COVID-19)(1,2). COVID-19 has swept the world since early 2020; at the time of writing this paper over 4.5 million deaths and 188 million infections have occurred worldwide, causing a global pandemic(3). There is currently no specific small molecule drug treatment for SARS-CoV, MERS-CoV, and SARS-CoV-2. Therefore, heavy priority has been placed on finding an effective antiviral drug, and several viral targets have been considered.

Two important viral targets controlling the production of functional proteins by SARS-CoV-2, a positive sense RNA virus, include Main Protease (Mpro) and Papain-like Protease (PLpro) encoded by the viral genome(4). A frameshift between Open Reading Frame 1a (ORF1a) and Open Reading Frame 1b (ORF1b) enables the production of two polypeptides: Polypeptide 1a (pp1a) and Polypeptide 1b (pp1ab). The viral proteases Mpro and Plpro process pp1a and pp1ab into 16 non-structural proteins necessary for viral assembly and replication as depicted in **Fig 1**. While inhibitors of individual proteases have been suggested(5,6), there are obvious benefits from the inhibition of *both* viral proteases in order to limit overall SARS-CoV-2 viral replication.

**Figure 1.**
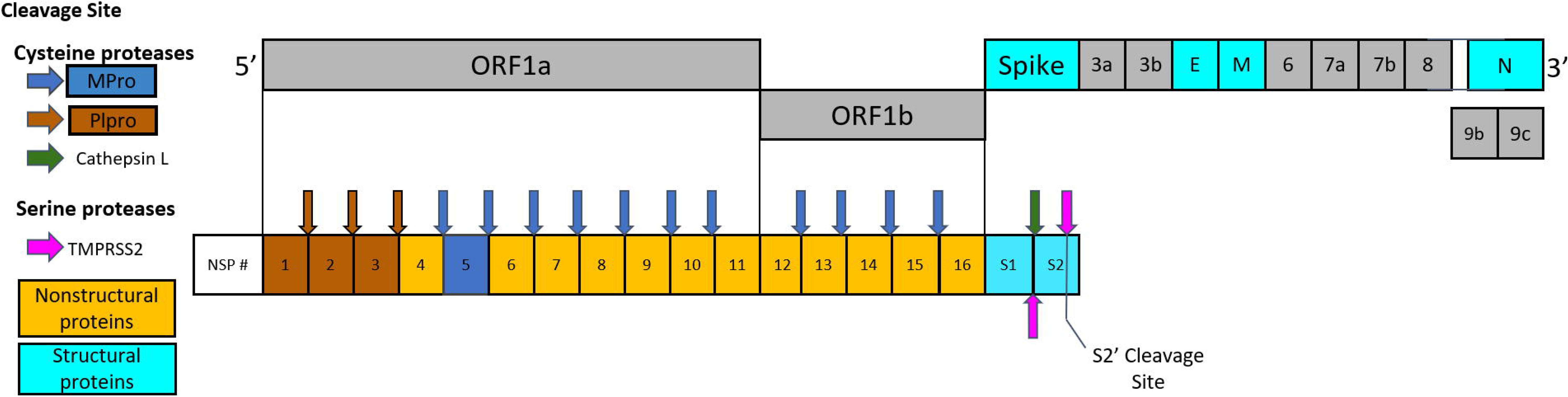
Schematic drawing of polypeptide expressed by SARS-CoV-2 including nonstructural proteins (NSP), structural proteins, and accessory proteins. Each represented viral protein has not been drawn to actual scale. Brown and blue arrows represent cleavage sites of SARS-CoV-2 Mpro and PLpro cleavage sites, respectively. Cathepsin L cleavage between S1/S2 subunits is represented by the green arrow. The cleavage sites of TMPRSS2 located between S1/S2 subunits and S’ cleavage site in S2 subunit were indicated by magenta arrows.

Moreover, the SARS-CoV-2 entry mechanism relies on two *human* proteases, the Transmembrane serine protease 2 (TMPRSS2) and endosomal cathepsin L to cleave the spike protein of SARS-CoV-2 which eventually binds to Angiotensin-converting enzyme 2 (ACE2), releasing the viral genome into the cytoplasm of the host cells and initiating the viral replication process(7–9). SARS-CoV-2 can exploit TMPRSS2 residing on cell membranes, facilitating their fusion. In addition to the TMPRSS2-mediated entry mechanism, SARS-CoV-2 can also utilize endosomal cathepsin L, an endosomal or lysosomal cysteine proteases(10). Cathepsin L was associated as a factor in COVID-19 compared to the uncorrelated lysosomal cysteine protease cathepsin B(11). These viral and human proteases are attractive targets for the design of anti-SARS-CoV-2 drugs. Our goal was to target *multiple* proteases that are necessary for the viral replication process with a *single* small molecule inhibitor. This approach is challenging but it may result in better antiviral drugs that are less sensitive to viral escape mutations resulting in drug resistance.

Several peptidomimetic compounds exhibiting effective inhibition against individual SARS-CoV-2 proteases have been identified(12–14) and many drugs have been investigated for drug repurposing for the COVID-19 treatment(15). However, our interests lie in covalent inhibitors working as multiple-target, anti-SARS-CoV-2 agents. One interesting compound is disulfiram which contains a thiuram disulfide covalent warhead at the center of the molecule. Disulfiram is a currently approved drug commonly prescribed as an aversive treatment of chronic alcoholism. Disulfiram primarily acts as an irreversible inhibitor of alcohol dehydrogenase by binding to a cysteine residue within the active site. The binding disrupts the breakdown of alcohol causing the development of resistance to alcohol consumption through minor toxicity and adverse effects occurring upon consumption(16).

Beyond the treatment of alcoholism, the distinct structure of disulfiram, containing the thiuram disulfide covalent warhead, is viewed as a key component to the drug’s activity in SARS-CoV-2 and is maintained as a point of interest in drugs targeting viral proteases. Previously, disulfiram was shown to inhibit both MERS-CoV and SARS-CoV PLpro with 14.6 μM and 24.1 μM IC_50_ values respectively. It was shown that disulfiram may act at the active site of the SARS-CoV PLpro, creating a covalent adduct with Cys112 that implied possible future uses of the drug in treatment against coronaviruses(17). More recently, disulfiram was characterized to display antiviral activity against multiple viral proteases including SARS-CoV-2 Mpro (IC_50_: 9.35 μM) and SARS CoV-2 PLpro (IC_50_: 6.9 μM) as well as several other viral cysteine proteases(18).

Furthermore, disulfiram has been characterized to block the pore formation of gasdermin D (GSDMD) through covalent targeting of cysteine 191, counteracting IL-β release and inflammation(19). In phosphoglycerate dehydrogenase (PHGDH) and caspase 1, disulfiram was covalently binds to cysteine residues which ultimately blocks the release of cytokines(20,21). Further characterized targets that have been shown to be inhibited by disulfiram are displayed in **Table 1**. With knowledge of this counteraction to inflammation and other favorable outcomes of disulfiram, it is currently in Phase 2 Clinical trials being identified as a potential therapeutic target for SARS CoV-2(22).

**Table 1-1.**
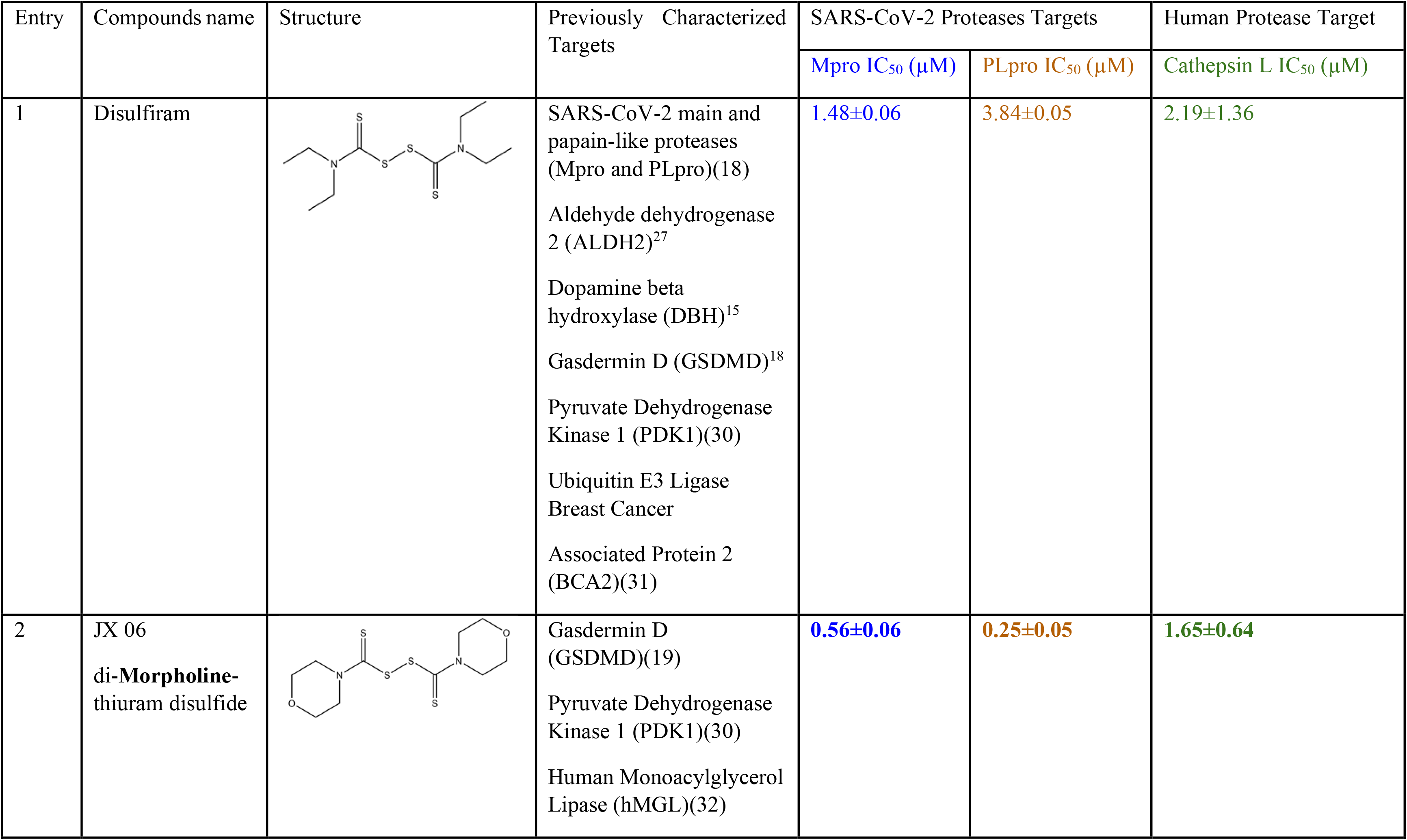
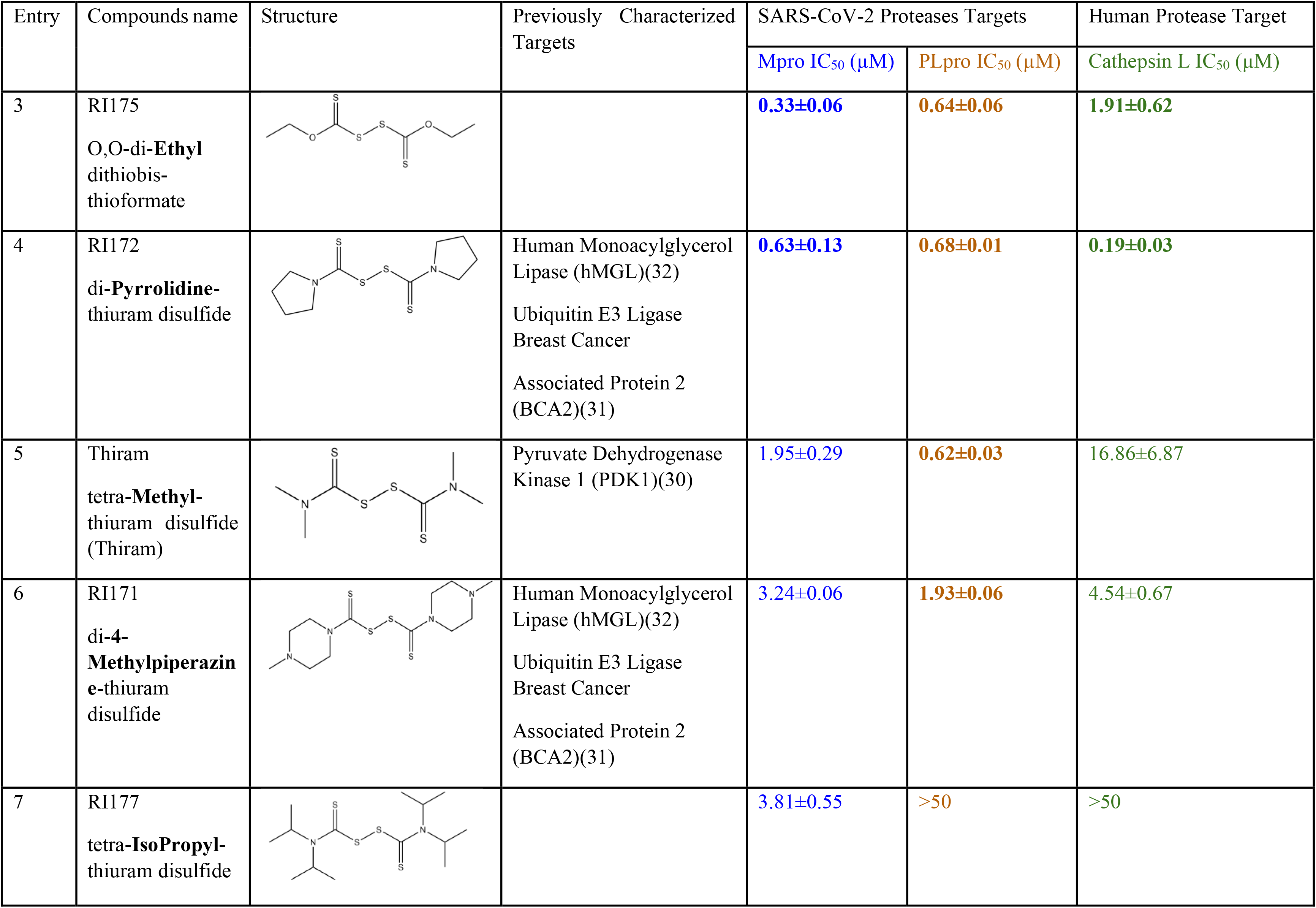
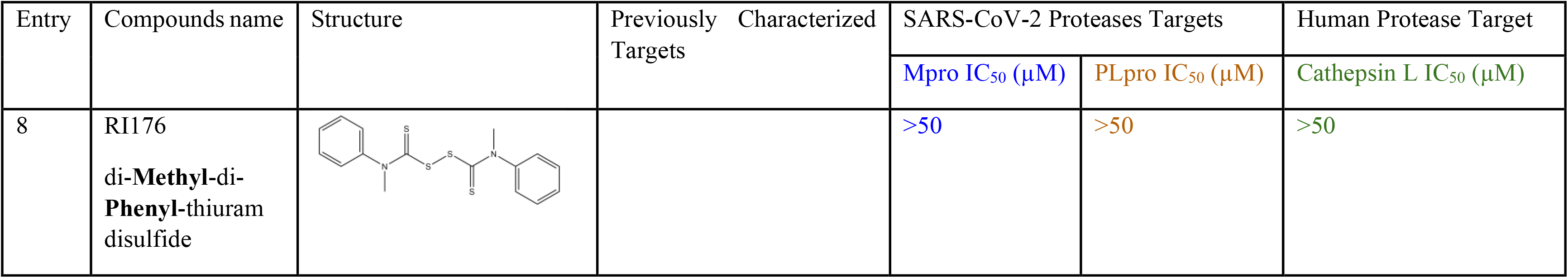
Structures of tested compounds with their known targets and enzymatic assays against SARS-CoV-2 Mpro and PLpro investigated in this study.

| Entry | Compounds name | Structure | Previously Characterized Targets | SARS-CoV-2 Proteases Targets |  | Human Protease Target |
| --- | --- | --- | --- | --- | --- | --- |
|  |  |  |  | Mpro IC<sub>50</sub> (μM) | PLpro IC<sub>50</sub> (μM) | Cathepsin L IC<sub>50</sub> (μM) |
| 1 | Disulfiram |  | SARS-CoV-2 main and papain-like proteases (Mpro and PLpro)(18)<br>Aldehyde dehydrogenase 2 (ALDH2)<sup>27</sup><br>Dopamine beta hydroxylase (DBH)<sup>15</sup><br>Gasdermin D (GSDMD)<sup>18</sup><br>Pyruvate Dehydrogenase Kinase 1 (PDK1)(30)<br>Ubiquitin E3 Ligase Breast Cancer Associated Protein 2 (BCA2)(31) | 1.48±0.06 | 3.84±0.05 | 2.19±1.36 |
| 2 | JX 06<br>di-Morpholine-thiuram disulfide |  | Gasdermin D (GSDMD)(19)<br>Pyruvate Dehydrogenase Kinase 1 (PDK1)(30)<br>Human Monoacylglycerol Lipase (hMGL)(32) | 0.56±0.06 | 0.25±0.05 | 1.65±0.64 |
|  |  |  |  | Mpro IC<sub>50</sub> (μM) | PLpro IC<sub>50</sub> (μM) | Cathepsin L IC<sub>50</sub> (μM) |
| 3 | RI175<br>O,O-di-**Ethyl**<br>dithiobis-<br>thioformate |  |  | **0.33±0.06** | **0.64±0.06** | **1.91±0.62** |
| 4 | RI172<br>di-**Pyrrolidine**-<br>thiuram disulfide |  | Human Monoacylglycerol<br>Lipase (hMGL)(32)<br>Ubiquitin E3 Ligase<br>Breast Cancer<br>Associated Protein 2<br>(BCA2)(31) | **0.63±0.13** | **0.68±0.01** | **0.19±0.03** |
| 5 | Thiram<br>tetra-**Methyl**-<br>thiuram disulfide<br>(Thiram) |  | Pyruvate Dehydrogenase<br>Kinase 1 (PDK1)(30) | **1.95±0.29** | **0.62±0.03** | **16.86±6.87** |
| 6 | RI171<br>di-**4-Methylpiperazin**<br>**e**-thiuram<br>disulfide |  | Human Monoacylglycerol<br>Lipase (hMGL)(32)<br>Ubiquitin E3 Ligase<br>Breast Cancer<br>Associated Protein 2<br>(BCA2)(31) | **3.24±0.06** | **1.93±0.06** | **4.54±0.67** |
| 7 | RI177<br>tetra-**IsoPropyl**-<br>thiuram disulfide |  |  | **3.81±0.55** | **>50** | **>50** |
|  |  |  |  | Mpro IC<sub>50</sub> (μM) | PLpro IC<sub>50</sub> (μM) | Cathepsin L IC<sub>50</sub> (μM) |
| 8 | RI176<br>di-**Methyl**-di-**Phenyl**-thiuram disulfide |  |  | >50 | >50 | >50 |

Analogs of disulfiram were tested on the SARS CoV-2 Papain Like and Main Protease to determine more effective and efficient inhibitors that take advantage of disulfiram’s mechanism of binding to the cysteine residue. To maintain this cysteine binding activity, our analogs contained the thiuram disulfide or dithiobis-(thioformate), varying in the functional groups beyond this center. Through experimental results, it was discovered that most analogs had either a similar or greater percent inhibition for both SARS-CoV-2 Mpro and PLpro in comparison to that of disulfiram. With overall relatively lower IC_50_ values, we attribute these analogs as viable triple target inhibitors of both SARS-CoV-2 proteases as well as human cathepsin L that may outperform disulfiram and warrant further testing and implementation in the clinical scene.

## Materials and methods

### Computational modeling

In our docking simulation, we obtained high resolution X-ray crystal structures of SARS-CoV-2 PLpro and Mpro from Protein Data Bank (PDB code: 5XBG and 6WX4, respectively) as the docking receptors. Protein-ligand complexes conformation and stability were determined by docking scores, which represent Gibbs free energy. The algorithm for sampling 3D conformation of ligands and pockets was generated randomly based on biased probability Monte Carlo. All docking simulations, scoring functions and pharmacokinetic properties were also predicted from ICM-Pro v3.9(23,24).

### Compounds and reagents

Disulfiram was purchased from Fisher Scientific (Hampton, NH). RI171 and RI172 were purchased from MolPort (Latvia) while JX 06 was purchased from Ambeed (Arlington Heights, IL). Thiram, RI175, RI176, and RI177 were obtained from Sigma-Aldrich (St. Louis, MO). All compounds were dissolved in dimethyl sulfoxide. All solvents were reagent grade, and all reagents were purchased from Sigma-Aldrich (St. Louis, MO).

### Recombinant protein and substrates

The recombinant SARS-CoV-2 main protease (Mpro) was expressed using the Mpro plasmid provided by Rolf Hilgenfeld(25) and purified as previously described(12,25). Recombinant proteases were purchased from following vendors: SARS-CoV-2 PLpro (Acro Biosystems, Newark, DE), Thrombin (R & D Systems, Minneapolis, MN), TMPRSS2 (Cusabio Technology LLC, Houston, TX), and human cathepsin L (R & D Systems, Minneapolis, MN). Protease substrates were purchased from following vendors: MCA-AVLQSGFR-K(DNP)-K-NH2 (R & D Systems, Minneapolis, MN), Z-RLRGG-AMC (Bachem Holding AG, Switzerland), Boc-VPR-AMC (Sigma-Aldrich, St. Louis, MO), Boc-QAR-AMC (Peptides International, Inc., Louisville, KY), and Z-FR-AMC (R & D Systems, Minneapolis, MN)

### Enzymatic inhibition assay

The protease enzymatic activities of SARS-CoV-2 Mpro and PLpro were measured using a FRET-based peptide substrate: MCA-AVLQSGFR-K(DNP)-K-NH2 and fluorogenic substrate: Z-RLRGG-AMC, respectively. The protease enzymatic activities of human thrombin, human TMPRSS2, and human cathepsin L were performed using fluorogenic substrates Boc-VPR-AMC, Boc-QAR-AMC, and Z-FR-AMC, respectively.

SARS-CoV-2 Mpro (50 nM final concentration) enzymatic reaction was carried out in reaction buffer containing 50 mM Tris-HCl pH 7.5, 150 mM NaCl, 1 mM EDTA and 0.01% Tween 20 using 10 μM of MCA-AVLQSGFR-K(DNP)-K-NH2 FRET based peptide as a substrate. SARS-CoV-2 PLpro (24.46 nM final concentration) enzymatic assays were performed in reaction buffer containing 50 mM HEPES pH 6.5, 150 mM NaCl and 0.01% Tween 20 using 50 μM of Z-RLRGG-AMC fluorogenic substrate. The positive control for these assays was 10 μM Ebselen. Human thrombin (678.3 nM final concentration) enzymatic assays were carried out in 50 mM Tris, 10 mM CaCl_2_, 150 mM NaCl, 0.05% Brij-35, pH 7.5 using 100 μM Boc-VPR-AMC as the substrate and 100 μM of dabigatran as positive control. Human TMPRSS2 (at 30 nM final concentration) was assayed in 50 mM Tris pH 8, 150 mM NaCl, and 0.01% Tween 20 buffer using 10 μM Boc-QAR-AMC substrate and 10 μM Nafamostat as positive control. Human cathepsin L (at 1 nM) activity assay was performed in 50 mM MES, 5 mM DTT, 1 mM EDTA, and 0.005% (w/v) Brij-35, pH 6.0 using 35 μM Z-FR-AMC as the substrate and 10 μM of E64 as positive control. The negative controls for all assays were 0.2% DMSO. All experiments were assayed in black 384-well microplate (BD Falcon) in the total volume of 50 μM at 37 °C. For Mpro FRET-based peptide substrate, the fluorescence signals were monitored at wavelengths of 320 and 400 nm for excitation and emission, respectively. Fluorescence signals of PLpro fluorogenic substrate were monitored at wavelengths of 360 and 460 nm for excitation and emission, respectively. All fluorescence signals were detected using Synergy HTX Multi-Mode Microplate Reader (BioTek) and data were visualized using Gen5 Software (BioTek). Dose response curves of each compound against all selected proteases were performed on 10 concentrations in triplicate ranging from 50 mM to 100 nM and IC_50_ values of each compound were calculated accordingly using SciPy and Matplotlib Python packages.

### Cells culture and immunofluorescence assay

For infectivity assay, Vero-E6 cells (2,000 cells/well, 384 well plate format) were used as host cells, infected with SARS-CoV-2 in an ‘multiplicity-of-infection’ value (MOI) of 1.0. Certain concentrations of compounds were spotted in the plates, followed by the Vero cells. Cells were incubated overnight at 37 °C with 5% of CO_2_. The cells were then infected by the virus and were incubated for 48h at 37°C with 5% CO_2_. The plates were then fixed with 4% PFA evaluated immunofluorescence signal for viral detection using Rabbit anti-nucleocapsid (GeneTex, cat# GTX135357) and anti-Rabbit Alexa488 as a secondary antibody. Cells were counterstained with DAPI. The image signals were analyzed using MetaXpress software to quantify individual cells and infected cells.

## Results

### SARS-CoV-2 Mpro and PLpro activity assay

Previous studies revealed that disulfiram has inhibitory activity against both Mpro and PLpro of SARS-CoV-2(18,26,27) and PLpro of SARS-CoV and MERS(16). In addition, disulfiram is currently being studied in phase two clinical trials as a therapeutic agent for SARS-CoV-2 infection(22). Disulfiram is known to form an irreversible covalent bond that targets active cysteine proteases(28). The excellent potency of disulfiram inspired us to further explore the inhibitory activities of compounds that contain thiuram disulfide or dithiobis-(thioformate), a necessary functional group for covalent binding to cysteine protease targets(29). Therefore, we selected six thiuram disulfide analogs with different N-substituents and one dithiobis-(thioformate), RI171 to RI177, to test against SARS-CoV-2 Mpro and PLpro. We performed fluorometric enzymatic assays to show the activities of Mpro and PLpro in the presence of selected compounds. The assays were performed using a FRET-based peptide substrate: MCA-AVLQSGFR-K(DNP)-K-NH2 at 10 μM and fluorogenic substrate: Z-RLRGG-AMC at 50 μM for Mpro and PLpro, respectively. As shown in **Fig 2** and **Table 1**, the inhibitory activities of RI172 and JX 06 against Mpro with the half-maximum inhibitory concentration (IC_50_) of 0.56 ± 0.06 and 0.63 ± 0.13 nM, respectively, showed approximately three folds improvement from disulfiram with an IC_50_ value of 1.48 ± 0.06 μM (2.1 ± 0.3 μM reported value(26), while Thiram, RI171, and RI177 showed slightly weaker inhibition effects compared to disulfiram. In addition, four of the thiuram disulfide compounds, JX 06, RI172, Thiram and RI171 showed significant potency improvement against PLpro with IC_50_ of 0.25 ± 0.05, 0.68 ± 0.01, 0.62 ± 0.03, and 1.93 ± 0.06 μM, respectively, compared to the reported IC_50_ of disulfiram, 4.11 μM (6.9 ± 4.2 μM reported value(18) especially for JX 06 which showed a 17-fold improvement over disulfiram.

**Figure 2.**
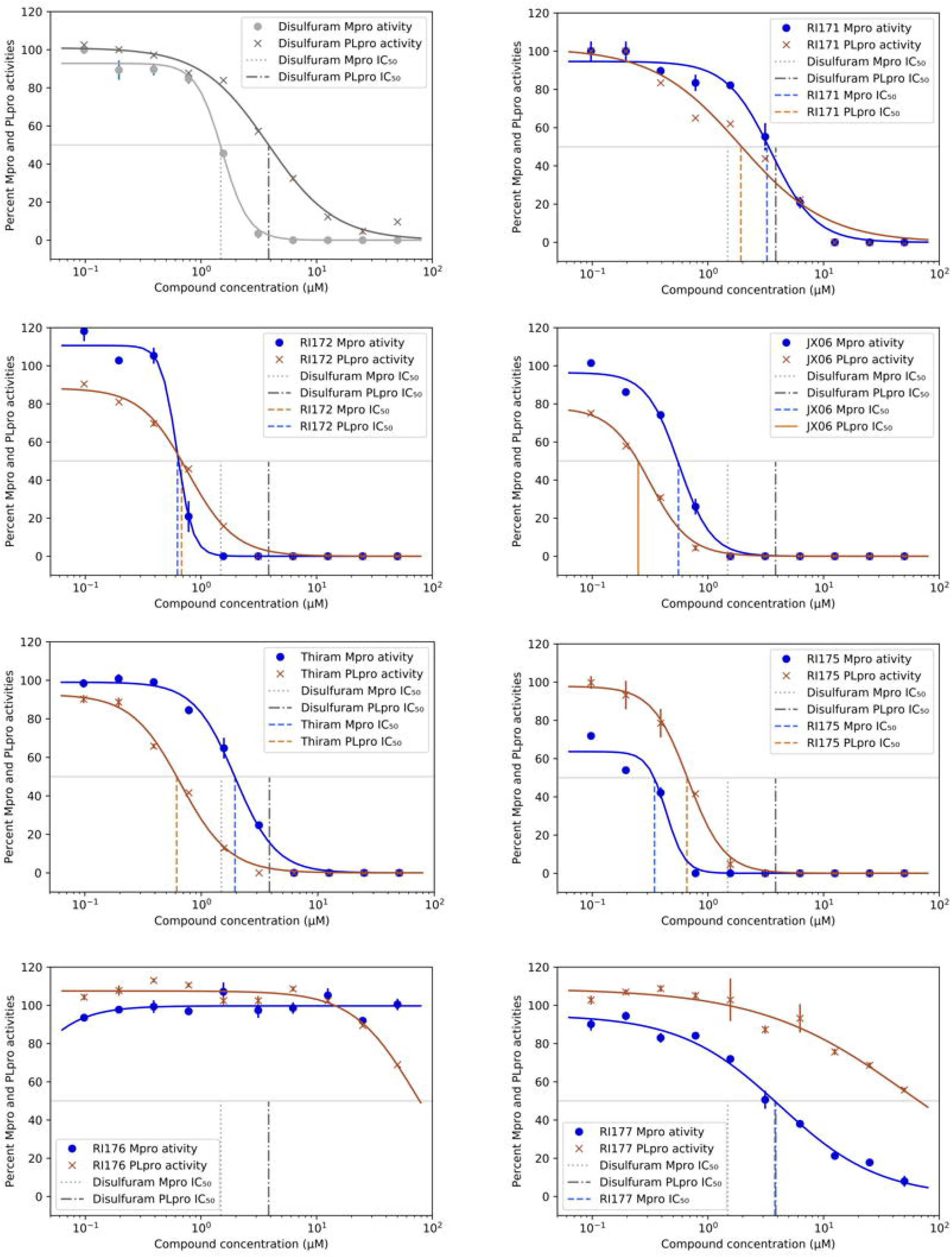
Selected thiuram disulfide or dithiobis-(thioformate) analogues show potent inhibition against SARS-CoV-2 Mpro and PLpro. Dose response curves of disulfiram, RI171, RI172, JX 06, Thiram, RI175, and RI177 against SARS-CoV-2 Mpro (blue) and PLpro (brown). Disulfiram dose response curve performed under the same conditions is shown for comparison. The proteolytic activities of both enzymes were determined by the enzymatic fluorescence assay and shown as percent activities relative to the negative control (DMSO). Error bars represent standard errors of independent three experiments.

These results suggest that thiuram disulfide is a necessary moiety to interact covalently with cysteine active residues of Mpro and PLpro. The replacement of ethylamine groups in disulfiram with morpholine, pyrrolidine, methylamine, and 4-methylpiperazine in JX 06, RI172, Thiram, and RI171, respectively, moderately enhance the potency of inhibitors against Mpro and PLpro, while substitution by isopropylamine in RI177 showed the preservation of inhibition of Mpro but completely lost the inhibitory activity (IC_50_ > 50 μM) against PLpro. Furthermore, the presence of N-methylaniline moiety in RI177 entirely lost the inhibition potency (IC_50_ > 50 μM) in both Mpro and PLpro. These results suggest that the substitution of bulky and flat moieties at terminal N-substituents are unfavorable for the improvement of inhibition against SARS-CoV-2 proteases for thiuram disulfide analogues. It is especially interesting that RI175 which contains dithiobis-(thioformate), unlike the rest of the compounds in this series, has shown potent inhibitory activity against both Mpro and PLpro, 0.33 ± 0.06 and 0.64 ± 0.06 μM, respectively.

### Cell entry proteins inhibition

Beside Mpro and PLpro, there are host proteases including TMPRSS2, cathepsin L, and furin that are necessary for SARS-CoV-2 viral replication. We selected both the endosomal cathepsin L and TMPRSS2 proteases that facilitate viral entry into the cell. SARS-CoV-2 binds to ACE2 on the cell surface and can enter cells either by single or dual activation mechanisms from Cathepsin L in the endosome and TMPRSS2 on the cell surface(10,33). In the previous section, several selected thiuram disulfide or dithiobis-(thioformate) analogues have shown excellent broadened inhibitory activity against both SARS-CoV-2 viral proteases. The mechanism of action of thiuram disulfide or dithiobis-(thioformate) analogues were known to bind to sulfhydryl groups of cysteine active residues of Mpro and PLpro becoming their oxidized cysteine form (**Fig 3**). Since cathepsin L contains cysteine as an active residue similar to Mpro and PLpro(34,35), we tested whether these compounds could also target cathepsin L, perhaps providing small molecules that target multiple key proteins for the viral replication process, producing a more favorable outcome for SARS-CoV-2 treatment. Thus, we also evaluated the inhibition activity of the selected compounds against 1 nM of cathepsin L using 30 μM of Z-Phe-Arg-AMC as a fluorogenic substrate; the dose response curves of each compound are shown in **Fig 4**. IC_50_ values of compounds JX 06 and RI175 (1.65 and 1.91 μM) have shown slight improvement from disulfiram (2.19±1.36 μM). RI172 showed excellent potency (0.19±0.03 μM) and significant enhancement, approximately 11.5 times more potent than disulfiram. Compounds RI171 and Thiram exhibited less potency compared to disulfiram, but they are still in acceptable range. On the other hand, thiuram disulfide analogs connected to bulky substituents isopropyl and methyl-di-phenyl, RI176 and RI177, respectively, did not seem to have any inhibitory effect to cathepsin L. This suggests that small alkyl or heterocyclic substituents on thiuram disulfide or dithiobis-(thioformate) analogs are more favorable than bulky substituents for strong inhibition to cathepsin L.

**Figure 3.**
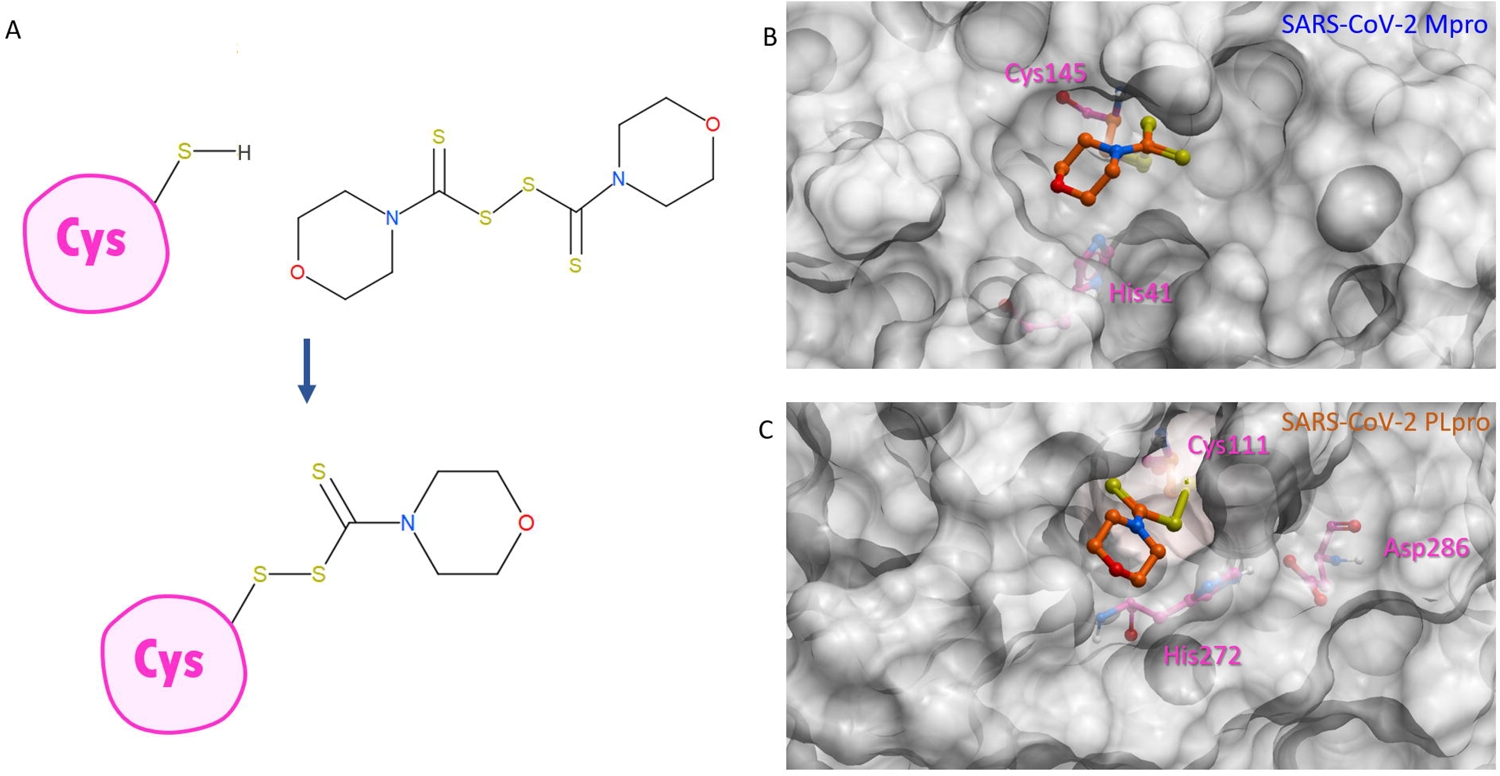
Binding conformation of thiuram disulfide or dithiobis-(thioformate). **A.** The mechanism of RI172 when it covalently reacts to active cysteine residue of SARS-CoV-2 Mpro and PLpro. **B.** Predicted 3D conformation of RI172 shown in orange stick in SARS-CoV-2 Mpro surrounded by catalytic dyads active residues Cys145 and His41. **C.** Predicted 3D conformation of RI172 shown in orange stick in SARS-CoV-2 PLpro surrounded by catalytic triads active residues Cys111, His272, and Asp286

**Figure 4.**
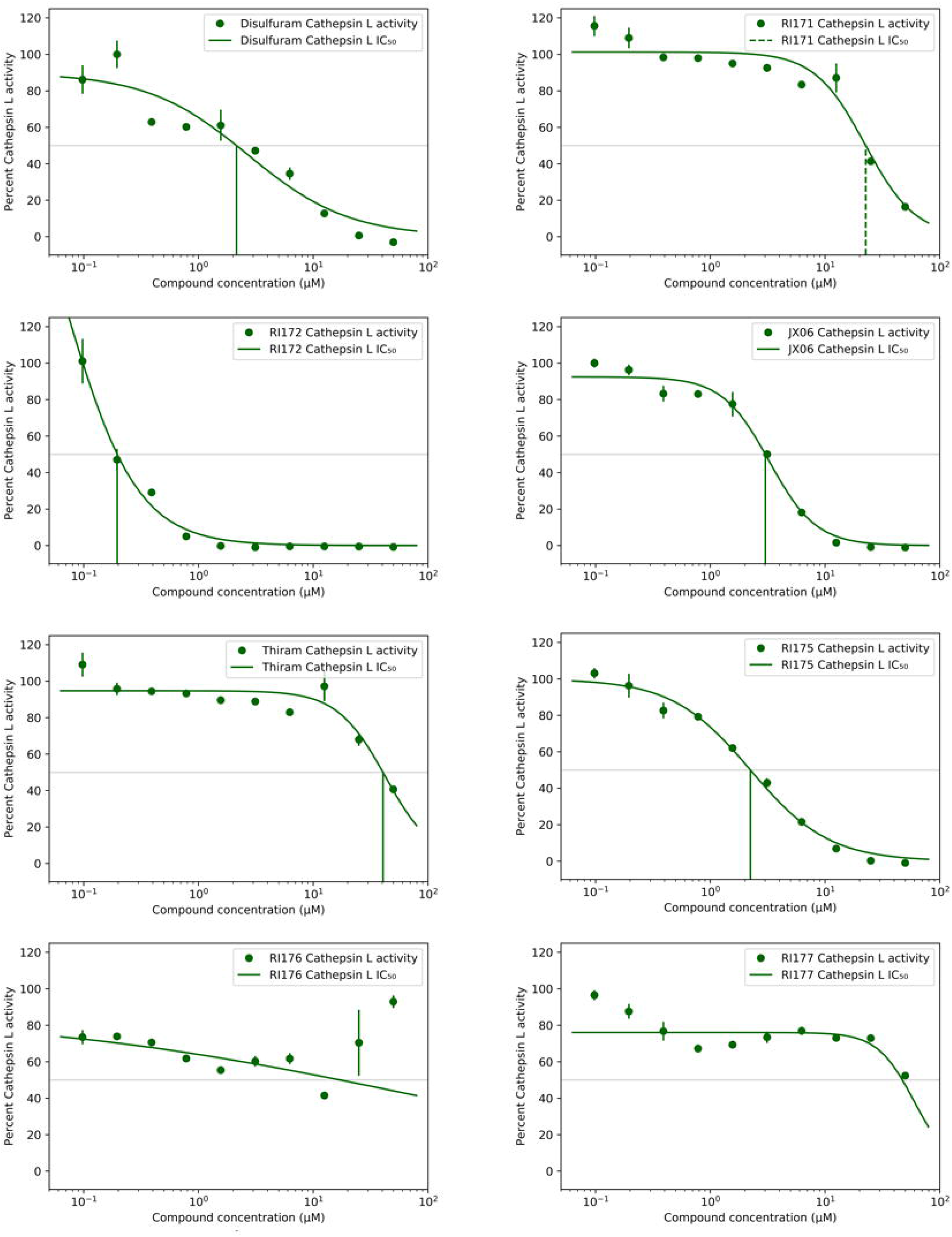
Thiuram disulfide or dithiobis-(thioformate) analogues inhibit human cathepsin L. Dose response curves of newly identified dual viral protease inhibitors disulfiram, RI171, RI172, JX 06, Thiram, RI175, RI176 and RI177 against human protease cathepsin L. Error bars represent standard errors of independent three experiments. The proteolytic activities of both enzymes were determined by the enzymatic fluorescence assay and shown as percent activities relative to the negative control (DMSO).

However, inactivation of cathepsin L does not completely prevent viral entry into the cell as SARS-CoV-2 utilizes TMPRSS2 as an alternative entry pathway(10). We tested our compounds against 30 nM of human TMPRSS2 using fluorogenic substrate 100 μM Boc-Gln-Ala-Arg-AMC, however there was no inhibitory activity of any compound up to100 μM (**Fig S1**). This proposes a cysteine protease specificity of the thiuram disulfide or dithiobis-(thioformate) analogues. According to these results, we have found triple inhibitors that target SARS-CoV-2 Mpro, SARS-CoV-2 PLpro, and human cathepsin L.

### Indirect effect in COVID-19 treatment through inhibition of Thrombin

Previously, it was shown that all thiuram disulfide or dithiobis-(thioformate) compounds have high potency against Mpro and PLpro of SARS-CoV-2 and human cathepsin L. Yet, we desired to further investigate if these compounds also inhibited other proteases related to the SARS-CoV-2 infection to provide further benefits in COVID-19 treatment. Severe pneumonia, acute inflammation and blood clots are common complications found in some severe COVID-19 infections(36–39). Coagulation is the usual immune response to bacterial or viral infections. However, COVID-19 is typically associated with hyper-inflammation activated by cytokines including interleukin-6 (IL-6), interleukin-1 (IL-1), and tumor necrosis factor-α (TNF-α) which can cause severe inflammation and harmful tissue damage if they were produced in excessive amount(40). Thrombin is a key protease for blood clot initialization by conversion coagulation factor from fibrinogen to fibrin. During inflammation, the anti-coagulation mechanism that prevents too much thrombin activation sometimes fails due to hyper-inflammation. This effect can increase risk of microthrombosis. Here, we tested if our selected compounds could slow down this hypercoagulation. However, when tested against 1 nM thrombin, even a 100 μM concentration of our analogs failed to show any significant inhibition (**Fig S1**).

### SARS-CoV-2 Infectivity Cells-Based Assay

Thiuram disulfide and dithiobis-(thioformate) inhibit three proteases that function in various critical stages of viral replication including viral entry and viral maturation. Targeting multiple proteases may have a therapeutic advantage over drugs that inhibit a single protease as it may reduce the emergence of drug resistant mutants. Therefore, we tested these triple-target compounds in a SARS-CoV-2 Vero E6 infection model. Cells were infected with SARS-CoV-2 in an MOI of 1.0 and treated with 1.4 μM, 6.0 μM and 24 μM of thiuram disulfide or dithiobis-(thioformate) for 48 h. We employed an immunofluorescence technique to visualize the number of viruses and host cells in the presence of these inhibitors. he half maximal effective concentration (EC_50_) and cytotoxic concentration (CC_50_) was determined (**Fig 5**). The relative antiviral activity was normalized to the untreated control (0% inhibition) and non-infected control (100% inhibition). The relative cell viability was calculated based on the number of host cells relative to the average number of untreated cells (100% cells viability). Consistent with the previous enzymatic activity results, JX 06, Thiram, RI171, RI172, RI175, and RI177 exhibited the EC_50_ in cells-based infectivity assay ranging from 2.5 to 12.6 μM. Our best compound from three protease *in vitro* assays, JX 06, showed the lowest EC_50_ at 2.5 μM which is approximately a *3.6*-*fold improvement compared to disulfiram*. Most compounds also exhibited acceptable range of therapeutic index values that allows the choice of appropriate concentrations for the treatment. These results provide the proof of concept of targeting multiple functional proteins involving the SARS-CoV-2 viral replication for the improvement of COVID-19 treatment.

**Figure 5.**
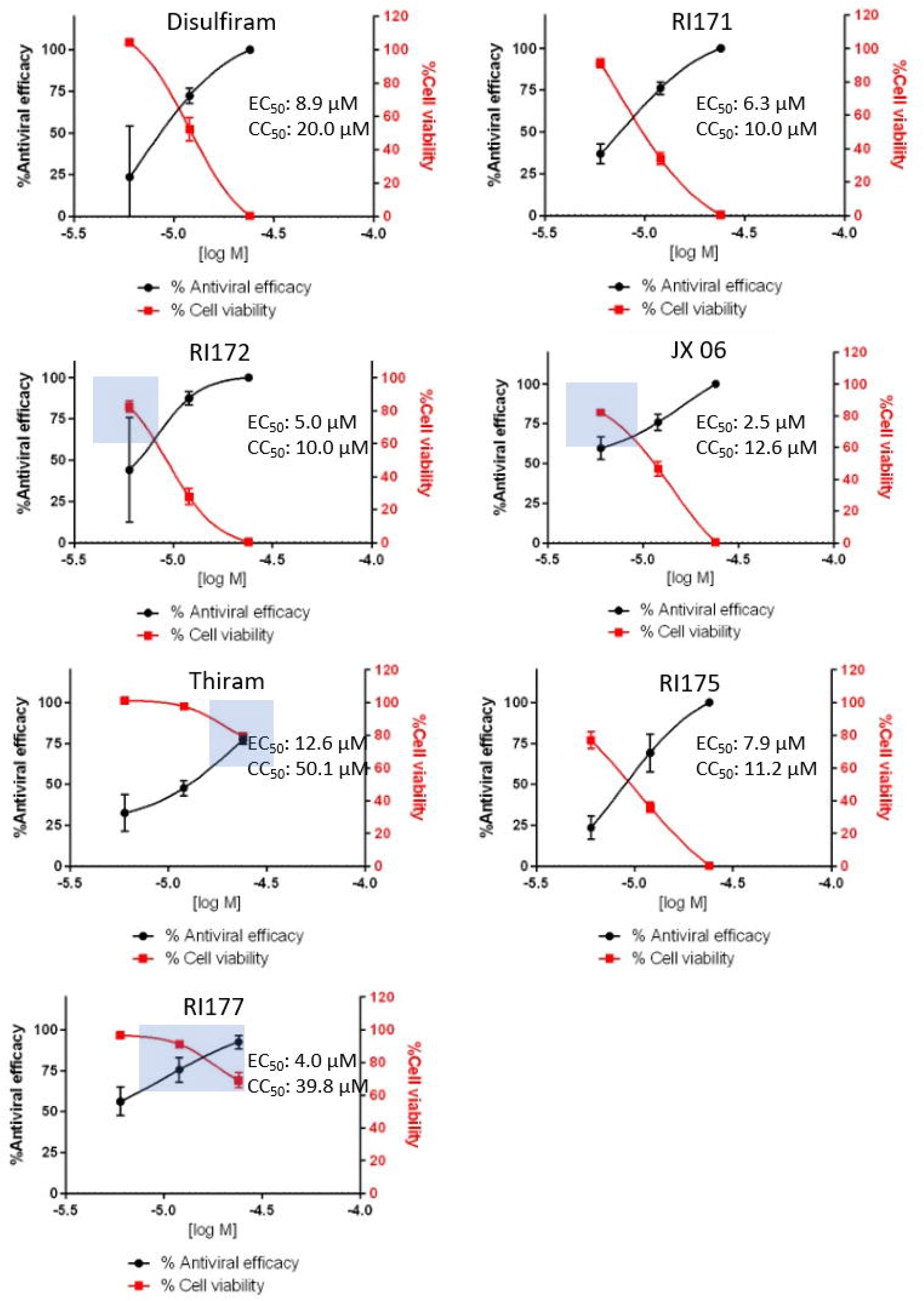
Selected thiuram disulfide or dithiobis-(thioformate) analogs reduce number of SARS-CoV-2 in certain concentrations. The antiviral activity (black solid circle) against SARS-CoV-2 and cell viability (solid red square) of disulfiram, RI171, RI172, JX 06, Thiram, RI175, and RI177 at 1.4, 6, 24 μM obtained from immunofluorescence signal and normalized compared to untreated control (0% inhibition) and non-infected control (100% inhibition and 100% cells viability). The half maximal effective concentration (EC_50_) and cytotoxic concentration (CC_50_) of each compound were evaluated from antiviral efficacy and cell viability normalized signals. Suggested therapeutic concentration range were shown in light blue area where the antiviral activity and cells viability are above 60%. Fitting was performed using GraphPad Prism software.

## Discussion

No effective therapy has been discovered for treatment of COVID-19 since the onset of the global pandemic. Treatment of COVID-19 has relied on available approved medicines such as α-interferon, the anti-HIV medication lopinavir, ritonavir, and remdesivir(41). More specific and effective therapies are necessary for post infection treatment. Most antiviral research is focused on specific single target inhibitors. On the other hand, targeting multiple proteins that function in viral replication can prevent the reduction of drug effectiveness caused by mutations of different target domains to develop drug resistance mutants. Disulfiram has been identified as a potent SARS-CoV-2 Mpro and PLpro inhibitor and is currently in phase two of clinical trials study for COVID-19 treatment. Moreover, disulfiram also covalently targets multiple enzymes including aldehyde dehydrogenase (ALDH2), dopamine beta hydroxylase (DBH), gasdermin D (GSDMD), pyruvate dehydrogenase kinase 1 (PDK1), ubiquitin E3 ligase breast cancer associated protein 2 (BCA2), human monoacylglycerol lipase (hMGL), and human cathepsin L(18–20,31,42). A multi-target pharmacological approach against enzymes that are directly and indirectly associated with SARS-CoV-2 infection may improve treatment of severe viral infection and reduce the emergence of drug-resistant variants. Here, we have shown that the five selected thiuram disulfide or dithiobis-(thioformate) analogs of disulfiram target key proteases in SARS-CoV-2 replication, namely the viral Mpro and PLpro enzymes in addition to human cathepsin L. We found that these compounds improve the inhibitory effectiveness against Mpro, PLpro, and cathepsin L 4.5, 17, and 11.5 times greater when compared to disulfiram, respectively. The cells-based infectivity assays also confirmed the efficacy of selected thiuram disulfide or dithiobis-(thioformate) compounds in viral infection treatment, especially for JX 06, our best compound, which showed 3.6 times more effectivity compared to disulfiram within the acceptable range of the therapeutic index. Although the suggested compounds could not cover all viral entry inhibition, a combination therapy with TMPRSS2 inhibitors such as nafamostat, camostat, and gabexate mesilate could improve the treatment efficacy in patients who have COVID-19. Beside targeting SARS-CoV-2 Mpro and PLpro, human cathepsin L, and alcohol dehydrogenase, disulfiram and JX 06, our best compound, is a known inhibitor of Pyruvate Dehydrogenase Kinase 1 (PDK1)(30) which functions as a switch from mitochondrial respiration to aerobic glycolysis. This process is called the “Warburg effect” and is thought to enhance malignancy^40,41^. SARS-CoV-2, similar to MERS-CoV, has a replication process that is promoted by the Warburg Effect via increasing the production of required resources for viral replication during the aerobic glycolysis enhancement(43,44). By blocking the activity of PDK1, JX 06 and all other thiuram disulfide or dithiobis-(thioformate) compounds in this series will help diminish SARS-CoV-2 viral replication, favorable for COVID-19 treatment. In addition, for most COVID-19 patients, there is extremely high production of proinflammatory cytokines that in excessive amounts can lead to acute lung injury and death^42,43^. Thiuram disulfide in disulfiram has been shown to covalently target Cys191 of Gasdermin D (GSDMD) and Cys116 of Phosphoglycerate dehydrogenase (PHDGH) which reduce the release of inflammatory cytokines(19,45). It can be assumed that dithiobis-thioformated analogues should have consistent effect in inhibition of GSDMD and PHDGH, like disulfiram. Treating COVID-19 patients with thiuram disulfide or dithiobis-(thioformate) analogues not only directly inhibits Mpro and PLpro viral proteins but could also benefit indirectly via disruption of viral replication processes and the reduction of excess inflammatory cytokines through PDK1, GSMD, and PHDGH inhibition. Even though there are concerns that disulfiram might be nonspecific towards cysteine proteases and might cause numerous side effects(18), the excellent improvement of the selected thiuram disulfide or dithiobis-(thioformate) compounds compared to disulfiram against the key proteins associated with COVID-19 suggest the potential for these drugs for oral administration in lower concentration against the virus.

## Conclusions

A single chemical substance may specifically inhibit both two viral proteases, Mpro and PLpro, and at least one human protease involved in viral infectivity cycle. This compound may also be relatively specific to the chosen viral and host targets. We identified the derivative of thiuram disulfide or dithiobis-(thioformate) which those favorable properties with the potential to slow down the viral infectivity cycle and modulate inflammatory responses.

## Acknowledgements

Authors would like to thank Elany Barbosa Da Silva, Ph.D. for her help in protease assay development, and Conall Sauvey for useful discussions and assistance. R.A. thanks NIGMS for the funding support R35 GM131881.

## Authors contributions

The research was designed and supervised by R.A., A.J.O., J.L.N-S., and J.H.M. The concept of hitting multiple proteins with the single compound was conceptualized by R.A. Disulfiram was proposed as the reference compound by R.A. Experiments were performed by I.M., D.S., P.F., M.A.G., and B.W. The manuscript was written by I.M., R.A., J.K., and J.Y.K.

## Supporting Information

**Figure S1. Screening on selected thiuram disulfide or dithiobis-(thioformate) compounds on TMPRSS2 and Thrombin.** Bar graph depicting percent inhibition of disulfiram, RI171, RI172, JX 06, Thiram, RI175, RI176 and RI177 at 0.1, 1, 10, 100 μM, against **(A)** 1 nM Thrombin and **(B)** 10 nM TMPRSS2 compared to known inhibitors E64 and Nafamostat, respectively. Error bars represent standard errors of independent three experiments. graphs were created using GraphPad Prism software.

