## Supplementary figures and images for "Discovery of Potent Triple Inhibitors of Both SARS-CoV-2 Proteases and Human Cathepsin L"

### S1_Figure.tif

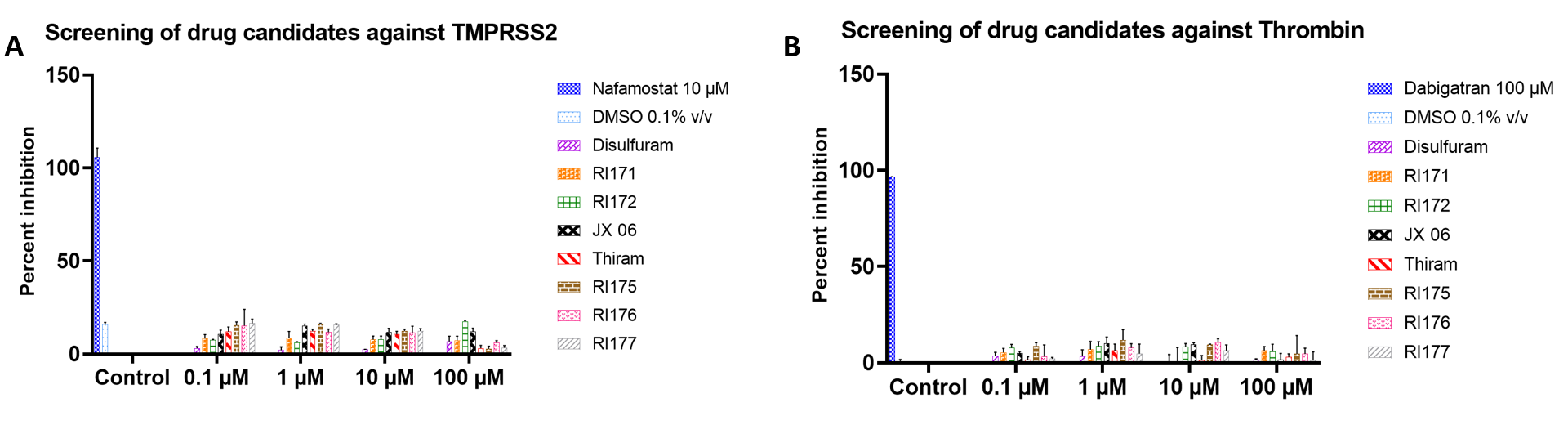
